# Niche-based deterministic assembly processes strengthen the effects of β-diversity on metacommunity functioning of marine bacterioplankton

**DOI:** 10.1101/2022.03.09.483723

**Authors:** Feng-Hsun Chang, Jinny Wu Yang, Ariana Chih-Hsien Liu, Hsiao-Pei Lu, Gwo Ching Gong, Fuh-Kwo Shiah, Chih-hao Hsieh

**Affiliations:** Institute of Oceanography, National Taiwan University, Taipei, Taiwan; Ecology and Evolutionary Biology, University of Michigan, Ann Arbor, USA; Department of Biotechnology and Bioindustry Sciences, National Cheng Kung University, Tainan, Taiwan; Institute of Marine Environment and Ecology, National Taiwan Ocean University, Keelung, Taiwan; Center of Excellence for the Oceans, National Taiwan Ocean University, Keelung, Taiwan; Research Center for Environmental Changes, Academia Sinica, Taipei, Taiwan; Institute of Ecology and Evolutionary Biology and Department of Life Science, National Taiwan University; National Center for Theoretical Sciences, Taipei, Taiwan

**Author notes:** Corresponding author: Feng-Hsun Chang, Or, Chih-hao Hsieh.

**Keywords:** Biodiversity-ecosystem functioning, Community assembly processes, Homogeneous versus heterogeneous selection, Predator-prey interaction, Metacommunity

## Abstract

Studies at a local community (α) level have shown that biodiversity positively affects ecosystem functioning due to niche-based deterministic processes like resource partitioning. Extending to a metacommunity (β) level, we hypothesize that β diversity also positively affects metacommunity functioning. We further hypothesize that the β diversity effect is stronger (more positive) when β diversity is increased by deterministic/non-random assembly processes. To test the hypotheses, we collected bacterioplankton along a transect of 6 stations in the southern East China Sea in 14 cruises. All 6 stations within a cruise were regarded as a metacommunity. For any pairs of the 6 stations, the Bray-Curtis index and summed bacterial biomass were calculated to represent β diversity and metacommunity functioning, respectively. We also calculated deviation of observed mean pairwise phylogenetic similarities among species from random to represent the deterministic influences of assembly processes. We found that bacterial β diversity generally positively affects metacommunity functioning; however, the β diversity effect varied among cruises. Cross-cruise comparison indicates that the β diversity effect increased with the deterministic assembly processes selecting for phylogenetically dissimilar species. This study extends the biodiversity-ecosystem functioning research to a metacommunity level, complementing the current framework by considering community assembly in natural environments.

## Introduction

Decades of studies have shown that, in a local community (α level), higher biodiversity increases ecosystem functions, such as resource use efficiency and biomass [1, 2]. This positive association has also been observed at the regional level (γ level) [3]. Whereas, interestingly at metacommunity (interconnected local communities) level, such a positive association between β biodiversity and ecosystem functioning can be expected because of the so-called “spatial insurance hypothesis” [4, 5]. In theory, the spatial insurance hypothesis asserts that different species sets, i.e., local communities, thrive in different local environments so that diverse local communities can collectively enhance the ecosystem functioning in a metacommunity. However, empirical studies do not always report positive effects of β diversity on the ecosystem functions of metacommunity [6]. Some studies show that β diversity is not important in predicting ecosystem functions [7]; a couple of recent studies even suggest that β diversity negatively affects ecosystem functions [8, 9].

Why is it that a positive association between β diversity and ecosystem functions has not been commonly observed in nature? This is a critical question in ecology, yet less studied compared to its α and γ level counterparts. Studying the relationship between β diversity and ecosystem functioning at metacommunity level is not merely a pattern-seeking exercise; rather, the attempt is to complement the existing framework of biodiversity-ecosystem functioning research. In the current biodiversity-ecosystem functioning framework, we have learned that selection effects and complementarity are the two main drivers underlying the positive effects of biodiversity [10, 11]. Selection effects occur when species favoring the ecosystem function are more likely to be included in a diverse community comparing to a species poor community [12–14]. Complementarity, on the other hand, may represent positive interactions and/or niche partitioning among the species comprising the community [15, 16]. However, these two drivers are not biological mechanisms, but properties emerging from assembly processes that determine which and how many species present in a community [17]. Since the assembly processes determine the species composition, diversity, and subsequently ecosystem functioning, these processes are worth considering to complement the biodiversity-ecosystem functioning research framework [6, 18].

In fact, community assembly processes are embedded in the calculation of β diversity because β diversity quantifies how species compositions are different from each other in a metacommunity [19, 20]. For example, some studies have statistically partitioned β diversity in order to understand whether species composition varies with certain environmental gradients [21, 22]. Others analyze β diversity by implementing multivariate statistical methods [23, 24], neutral-theory-based process models [25, 26], or null model methods [27–29] in order to infer how the species compositions are determined. By investigating the relationship between β diversity and functions of metacommunity, the influences of assembly processes should be inherently taken into account.

The assembly processes determining the species compositions and β diversity of a metacommunity can be stochastic or deterministic [30, 31]. The stochastic processes mean that random chance events govern species’ birth, death, presence/absence, and thus population size of species [26, 32]. Either due to random environmental fluctuations (i.e. environmental stochasticity) or independent to environments (i.e. demographic stochasticity), species’ random birth, death, and dispersal can cause the species compositions to be different [33, 34]. On the contrary, the deterministic processes are non-random and niche-based processes, representing biotic (e.g. species competition) or abiotic interactions (e.g. habitat filtering) among species [35, 36]. The stochastic and deterministic processes collectively determine the β diversity of a metacommunity [27, 37].

More importantly, not only do stochastic and deterministic assembly processes determine β diversity, the assembly processes also determine whether β diversity enhances, decreases, or exerts no clear influences on the functions of metacommunity [38, 39]. For example, some species can persist in a metacommunity and contribute to the β diversity due to stochastic processes like dispersal, but these species are not able to utilize resources and perform ecological functions [32, 40]; under such circumstance, β diversity does not necessarily enhance the functions of metacommunity. By contrast, deterministic assembly processes selecting species that can capture resources and survive should result in positive β diversity effects on metacommunity functioning. We argue that when the metacommunity is governed more by deterministic and less by stochastic assembly processes, β diversity would have stronger effects on ecosystem functions.

To evaluate whether a metacommunity is governed by stochastic or deterministic assembly processes, we resort to the degree of relatedness, i.e., phylogenetic similarity, among species [41, 42] because phylogenetic similarity is considered as an imprint left by evolutionary and ecological processes [43, 44]. We then apply the null model approach on the phylogenetic similarity among species in a metacommunity to infer the relative importance of stochastic versus deterministic assembly processes [27]. The null model approach is elaborated in the method section. In short, the null distribution of phylogenetic similarity is derived by randomization to represent stochastic processes. When phylogenetically similar species are selected by homogenizing deterministic processes, the observed phylogenetic similarity will be negatively deviated from random. In contrast, when diversifying deterministic assembly processes select for phylogenetically dissimilar species and increase β diversity, the observed phylogenetic similarity is expected to be positively deviated from random.

To investigate the influences of community assembly processes on the association between bacterial β diversity and metacommunity functioning, we collected bacterial samples along a transect of 6 stations in the southern East China Sea (ECS) in 14 cruises (map in Figure 1A). The southern ECS is a hydrographically complex region [45, 46]. The inner shelf is dominated by river inputs and coastal currents that are typically non-saline and rich in terrestrial nutrients [47, 48]. In the middle shelf, the Taiwan Warm Current (Taiwan Strait Current) is warm and nutrient-depleted in the surface but saline and nutrient-rich in the bottom layer [47, 48]. To the northeast of Taiwan at the edge of continental shelf, the upwelled subsurface water from the Kuroshio Current bring ample amount of saline and nutrient-rich water to the ECS [49]. Interactions among these water masses vary spatially and seasonally to determine the dynamics of nutrients, chlorophyll-a, primary production [50] and bacterial biomass and production [51, 52]. These spatial and temporal complexities also create among-cruise variations in deterministic versus stochastic assembly processes for bacterial communities in the southern ECS [53].

**Figure 1.**
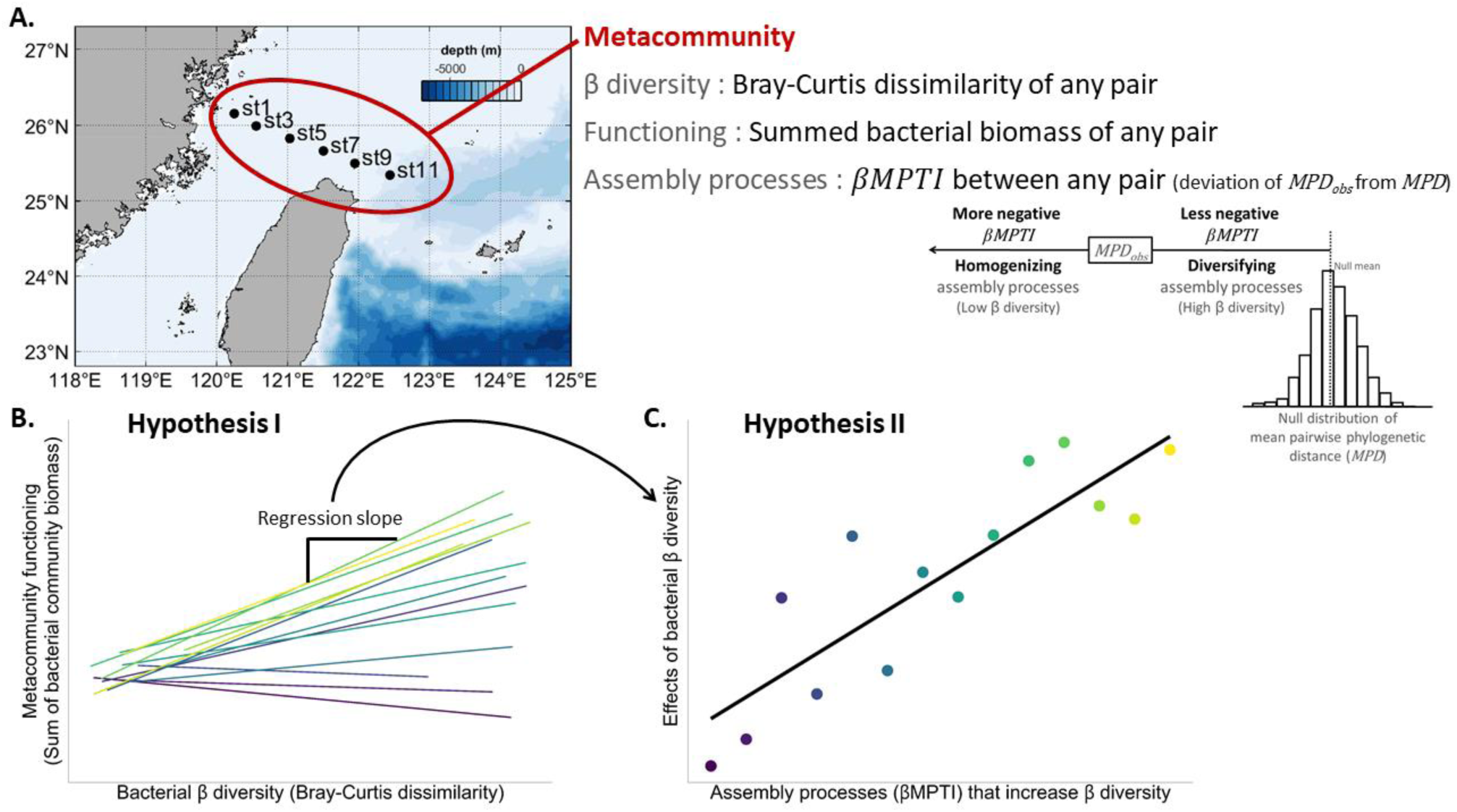
Schematic plots visualizing the sampling sites, analytical flow and hypotheses. Panel A illustrates the 6 sampling stations in the southern East China Sea, as well as the indices of β diversity, functioning, and assembly processes of the metacommunity; the inset panel describes the interpretation of the index of assembly processes (*βMPTI*). Panel B shows the expectation of Hypothesis I: Bacterial β diversity increases the sum of bacterial community biomass. Each colored line in panel B represents the linear model that regresses metacommunity functioning on bacterial β diversity for each cruise, and the regression slope is an indicator of β diversity effect. The regression slopes are shown as the colored dots in panel C, which shows the expectation of Hypothesis II (panel C): The effect of bacterial β diversity on the sum of bacterial community biomass is stronger when the β diversity effect is positively associated with deterministic assembly processes.

In this study, we defined a bacterial metacommunity as the 6 stations within each of the 14 sampling cruises. First, we test Hypothesis I: Bacterial β diversity increases the summed bacterial community biomass (Figure 1B). Specifically, for any pair of these 6 stations, we calculated the Bray-Curtis index as the bacterial β diversity and the sum of bacterial community biomass as the indicator of metacommunity functioning. Then, the effect of β diversity on metacommunity functioning could thus be estimated as the slope when regressing metacommunity biomass against the β diversity for each bacterial metacommunity (i.e., each cruise) (Figure 1B). Second, we test Hypothesis II: Based on cross-cruise comparison, the bacterial β diversity effect is stronger (more positive) when β diversity is increased by deterministic assembly processes that increase β diversity (Figure 1C). That is, we expect that the effects of bacterial β diversity (defined as the estimated slope as explained in Figure 1B) would be more positive when the observed phylogenetic similarity is positively deviated from random.

## Methods

### Bacteria biomass, community composition and environmental variables

We collected bacteria from a transect of 6 stations in the southern ECS (map of Figure 1A). We visited this transect in 14 cruises from April 2014 to July 2017, so that a total of 84 samples were obtained. At each sampling station, GoFlo bottles (General Oceanics) mounted on a conductivity, temperature and depth profiler (CTD profiler, Sea-Bird Electronics, Bellevue, WA, United States) were used to collect ∼20 L of seawater from 5 m beneath surface. The seawater was pre-filtered through a screen mesh with 20 μm openings to remove large particles like zooplankton. With the pre-filtered seawater, we estimated the bacterial biomass with a flow cytometer and revealed the taxonomic composition with high throughput sequencing techniques.

To estimate the bacterial community biomass, we preserved 10 mL of pre- filtered seawater in glutaraldehyde with final concentration equaled to 0.1%. The samples were frozen with liquid nitrogen and stored at −80°C before counting with a flow cytometer. Prior to processing with a flow cytometer, the samples were stained with SYBR Green I and then incubated for 15 minutes at room temperature in dark. We counted bacterial density with the CyFlow® Space (PARTEC) at a rate allowing <1,000 events per second to avoid particle coincidence. Bacterial density was then used to estimate bacterial community biomass with a carbon conversion factor of 2 × 10^-14^ gC/cell [54]. To account for the potential predators of bacteria, we counted the heterotrophic nanoflagelltes (HNF) under a fluorescence microscope [55].

Specifically, we additionally preserved 50 mL of pre-filtered seawater in neutralized formalin with final concentration equaled to 2% and stored the samples at 4°C. Samples were identified and counted using an inverted epifluorescence microscope (Nikon-Tmd 300) at 200× or 400× magnification [56]. HNF community biomass was finally converted using a carbon conversion factor of 4.7 × 10^-12^ gC/cell [57].

The rest of the pre-filtered seawater was filtered sequentially through a 1.2 μm and a 0.2 μm-pore-size filter (Millipore Isopore^TM^ hydrophilic polycarbonate membrane). The 0.2 μm-pore-size filter was frozen immediately with liquid nitrogen and then stored at −80℃ before molecular analysis [58]. From the filter, we extracted and sequenced DNA and processed sequences for bacteria composition. The methods of DNA extraction and sequence processing were briefly introduced here. Detailed methods were explained in Supplement 1. Total DNA was extracted from the 0.2 μm- pore size filter with the PowerWater DNA Extraction Kit (PowerWater, Qiagen) according to the manufacturer’s instructions. DNA extracts from the filters were used as templates of polymerase chain reaction (PCR) to amplify the 16S rDNA for bacterial community. PCR was performed in two steps to gain better reproducibility and consistent results [59]; see Supplement 1 for details of the two-step PCR procedures. After obtaining 16S rDNA sequences, the DADA2 pipeline was used for quality filtering and assembling sequences into amplicon sequence variant (ASV) [60]; see Supplement 1 for sequence merging procedures. Taxonomy assignment was performed on ASVs to recognize and select for those classified under the bacteria kingdom based on the Silva 132 database [61]. Finally, in order to obtain the phylogeny, we used maximum likelihood method (with negative edges length = 0) [60] to build phylogenetic trees for bacteria from 16S rDNA of all cruises.

For each sampling station, environmental variables, including temperature, salinity and photosynthetic active radiation, were recorded by the CTD profiler. In addition, 100 ml of seawater samples for chlorophyll-a, total dissolved inorganic nitrogen and phosphate concentrations, were collected and measured according to the standard methods [50].

### β diversity and bacterial biomass

In this study, we defined a bacterial metacommunity as the communities collected from the 6 stations within each of the 14 sampling cruises. For any pair of these 6 stations (15 pairs in total), we calculated the summed bacterial community biomass as the indicator of metacommunity functioning and the bacterial β diversity as the Bray-Curtis dissimilarity index. Before calculating the β diversity of bacteria, each community was resampled once to achieve the same number of reads, i.e., 13 129, the minimum reads among stations across cruises. Although this procedure addressed the disparity issue (i.e., unequal reads among stations), it is still not appropriate to compare the relative abundance of AVSs across stations [62].

Therefore, we applied Chao et al’s method to rarefy bacteria communities in order to have a fair among-station comparison [63–65]. With the rarefied compositions, we calculated the Bray-Curtis dissimilarity for any pair of the 6 stations as the β diversity of bacterial metacommunity for each cruise.

### Community assembly processes

To infer the community assembly processes governing the bacterial metacommunity, we calculated the β mean pairwise taxonomic index (*βMPTI*). The *βMPTI* extends Webb’s net relatedness index [41, 66] from within a community at α level to across two communities at β, i.e. metacommunity level [67]. The *βMPTI* quantifies the deviation of the observed mean pairwise phylogenetic distance (*MPD_obs_*) in two communities from a null distribution of mean pairwise phylogenetic distance (*MPD*) in the same two communities. The *MPD_obs_* was calculated as the mean branch length among all pairs of species in the two communities comprising the metacommunity. The null distribution of *MPD* is generated by random processes so that the distribution represents the values of *MPD* if a metacommunity is assembled completely by random processes. The deviation of *MPD_obs_* from the null thus represents the non-random/deterministic processes by which a metacommunity is assembled. To generate a null distribution of *MPD* by random processes, we first randomly shuffled the phylogeny (i.e., randomly shuffled the tips of the phylogenetic tree) of all species in a metacommunity (i.e., any pairs of the 6 stations within a cruise). With the randomized phylogeny, we calculated *MPD* of each metacommunity. Repeating this randomization technique 999 times, we generated the null distribution of *MPD*. Finally, *βMPTI* = (*MPD_obs_* − *meanMPD_null_*)/*sdMPD_null_*, where *meanMPD_null_* represents the mean of the null distribution of *MPD*, and *sdMPD_null_* represents the standard deviation of the null distribution of *MPD*.

The *βMPTI* is nearly identical to the β net relatedness index (*βNTI*). The only difference is that we multiply *βNTI* by -1 so that the sign of *βMPTI* intuitively represents whether the observed phylogenetic distance is more similar (more negative) or less similar (less negative) from random. Negative *βMPTI* means that species in the two communities are more phylogenetically similar than expected from random. More negative *βMPTI* thus implies that the metacommunity is subject to assembly processes selecting for species that are phylogenetically more similar to each other. Following the same logic, less negative *βMPTI* hints at the assembly processes that select for phylogenetically less similar (i.e., more dissimilar) species.

### Biodiversity-ecosystem functioning relationships and impacts of community assembly processes

With the bacterial β diversity and biomass, we tested Hypothesis I: Bacterial β diversity and the summed community biomass is positively associated (Figure 1B). To do so, we performed generalized linear mixed effect model (GLMM) to regress the sum of bacterial community biomass against β diversity, with cruise as the random effect. The sum of bacterial community biomass and β diversity were log transformed to improve normality. Making cruise as a random effect should account for the temporal autocorrelations among cruises. To statistically account for potential confounding factors, we conducted backward selection by first including all environmental variables and step-wisely removing the variables that were not significant based on p-values. The environmental variables include the average temperature, salinity, total dissolved inorganic nitrogen, phosphate, photosynthetically active radiation (PAR), and chlorophyll-a concentration (collected at the same depth as the bacteria samples) of the two stations comprising the metacommunity.

Next, we proceeded to test Hypothesis II: The effect of bacterial β diversity on the sum of bacterial community biomass is stronger when β diversity is increased by niche-based, i.e., non-random/deterministic processes (Figure 1C). First, we regressed the bacterial β diversity against *βMPTI*, in order to check whether in general β diversity increased with assembly processes that selected for phylogenetically dissimilar species. Next, the effect of β diversity was estimated from the generalized linear model (GLM) that regressed the sum of bacterial community biomass on β diversity of any pair of the 6 stations for each cruise. Here, the resulting 14 regression coefficients (i.e., slopes) were regarded as the effects of β diversity on metacommunity functioning. We finally regressed these regression coefficients against the mean *βMPTI* of each sampling cruise, to test if the effects of β diversity would increase with the assembly processes that selected for phylogenetically dissimilar species. Similarly, we also conducted backward selection to statistically account for potential confounding environmental variables.

### Computation

We used the “phyloseq” package to perform sequence subsampling to achieve parity in total number of reads [68], the “iNEXT” package to perform rarefaction [65], the “phangorn” package to build phylogenetic trees [60], the “picante” package to calculate phylogenetic distances to derive *βMPTI* [69] and the “nlme” package to perform generalized linear mixed effect models (GLMM) [70]. All packages were built and computation was carried out in R ver. 4.1.1 [71]

## Results

In the southern East China Sea (ECS), physical and chemical environments significantly changed from the inner to outer shelf. At the most inner shelf station, temperature and salinity were significantly lower than other stations, especially in the springtime (p < 0.01; Figure S2-1). This signaled the river runoff of the Min River that significantly increased nitrogen and phosphorous concentrations and caused the chlorophyll a concentration to be significantly higher at the most inner shelf station than other stations (p < 0.01; Figure S2-1), except for the station northeast of Taiwan (station 9; Figure S2-1). The station at the northeast of Taiwan is impacted by the all year round upwelling of subsurface Kuroshio waters, so that it is generally nutrient-rich [72]. However, the nutrients supplied by upwelling were lower than the river runoff, so that bacterial community biomass at this station was still significantly lower than the inner shelf stations (Station 1 and 3; p < 0.01; Figure S2-1).

Associating with the environmental variations, the bacterial community also showed a clear compositional shift from the inner to outer shelf of southern ECS (Figure S2-2). Specifically, the bacterial communities at the inner shelf were more positively associated with temperature and salinity, while the communities were more positively associated with total inorganic nitrogen and phosphorous at the outer shelf (Figure S2-2). These results suggest that the environment is indeed spatially heterogeneous, and such spatial heterogeneity in turn affects the community composition and biomass of bacterioplankton in the southern ECS.

For Hypothesis I, we found that the sum of bacterial community biomass was positively associated with bacterial β diversity (univariate regression coefficient = 0.17; Figure 2), even after accounting for potential confounding environmental variables (Table 1). Our results thus support Hypothesis I to some extent. However, this association was highly variable, judging from the low marginal R^2^ (0.01) of the GLMM. When fitting generalized linear model for each cruise, 8 out of 14 regression slopes were non-significant (colored dash lines in Figure 2), indicating that the effect of β diversity was not significant in these 8 bacterial metacommunities. In addition, we found that *βMPTI* varied significantly among the 14 sampling cruises and seemed to be lower in those 8 cruises (Figure S3-1). These results suggest that the effect of bacterial β diversity on the sum of bacterial communities is generally positive, but it could vary with the assembly processes of metacommunity.

**Figure 2.**
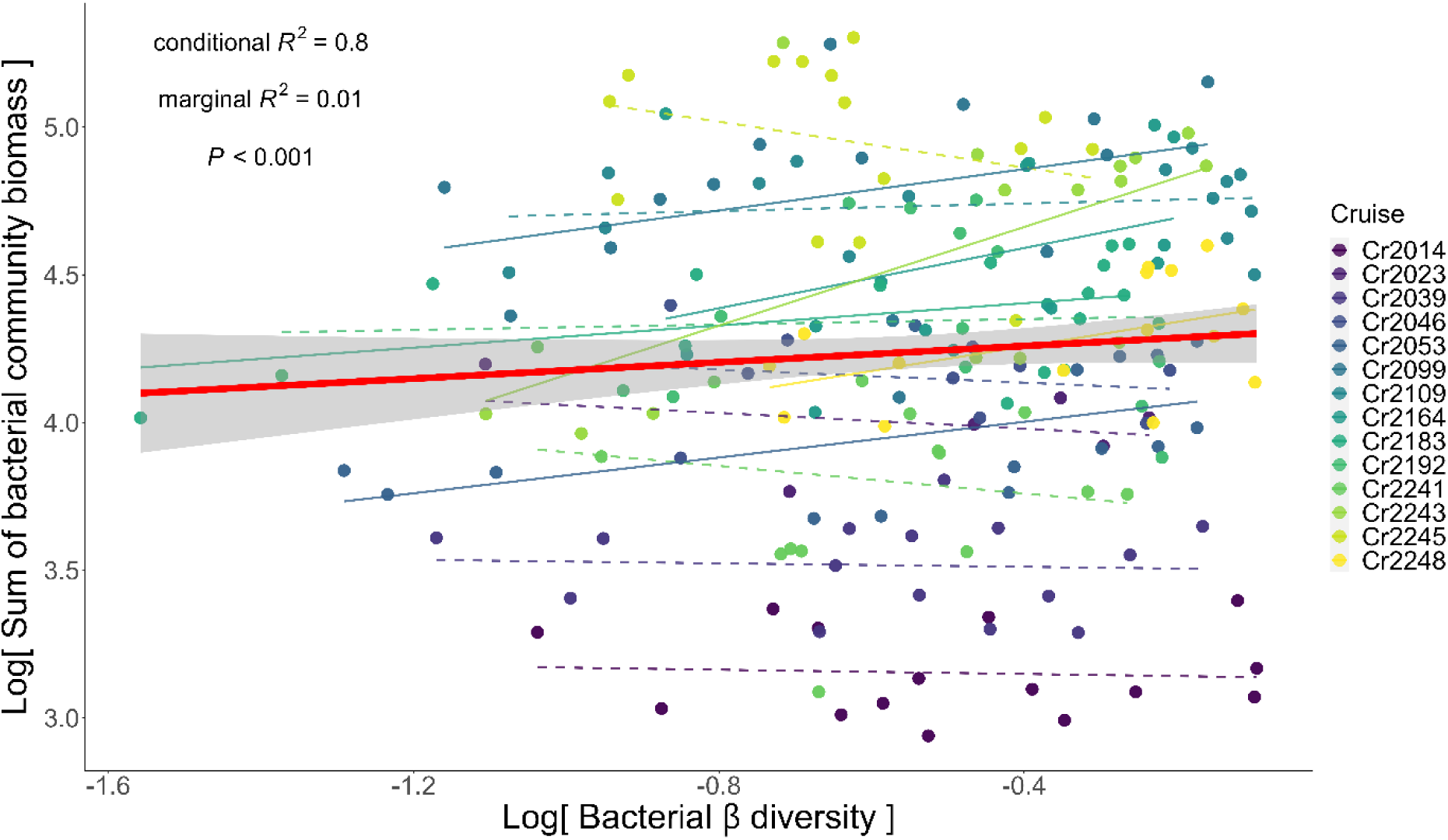
The association between the sum of bacterial community biomass and β diversity. The bold red solid line indicates the regression line fitted by the generalized linear mixed effect model (GLMM), with cruise as the random effect. Grey shaded area represents the confidence interval of regression. The colored dots represent measurements from different sampling cruises. The thin colored line represents the regression line fitted by generalized linear model regressing the sum of pair-wise bacterial community biomass on β diversity for each cruise. These regression lines vary substantially among cruises. Colored solid and dashed lines represent significant and non-significant regression slopes, respectively.

**Table 1.** Results of generalized linear mixed effect model (GLMM) testing Hypothesis I: Bacterial Bray-Cutis dissimilarity index increases the sum of bacterial community biomass (Figure 1A). The model presented here is the results after backward stepwise selection to statistically control for potential confounding factors. Steps of the backward stepwise selection is listed in Supplementary Table S4-1. The results indicate that, after accounting for environmental variables, bacterial β diversity positively affected metacommunity biomass. This conclusion is qualitatively the same as that drawn from the univariate GLMM (Figure 2).

| Independent variable | Regression coefficient | Standard error | p-value |
| --- | --- | --- | --- |
| Log (Bacterial $\beta$ diversity) | 0.09 | 0.04 | 0.04 |
| Log (Salinity) | -3.86 | 0.44 | <0.01 |
| Log (Total inorganic nitrogen) | -0.02 | 0.01 | <0.01 |

We then test Hypothesis II: When β diversity is increased by niche-based, i.e., non-random/deterministic processes, the effect of bacterial β diversity on the sum of bacterial communities is more positive (Figure 1C). We first noticed that *βMPTI* was generally negative in our study (mean = -1.22; standard deviation = 0.76), which indicated that the bacterial metacommunity in the southern ECS was generally subject to homogenizing assembly processes. However, when *βMPTI* became less negative, bacterial β diversity was also higher (univariate regression slope = 0.09 and p-value < 0.01; Figure 3). These results suggest that β diversity of metacommunity is increased when deterministic assembly processes weaken the influences of homogenization by selecting for phylogenetically dissimilar species.

**Figure 3.**
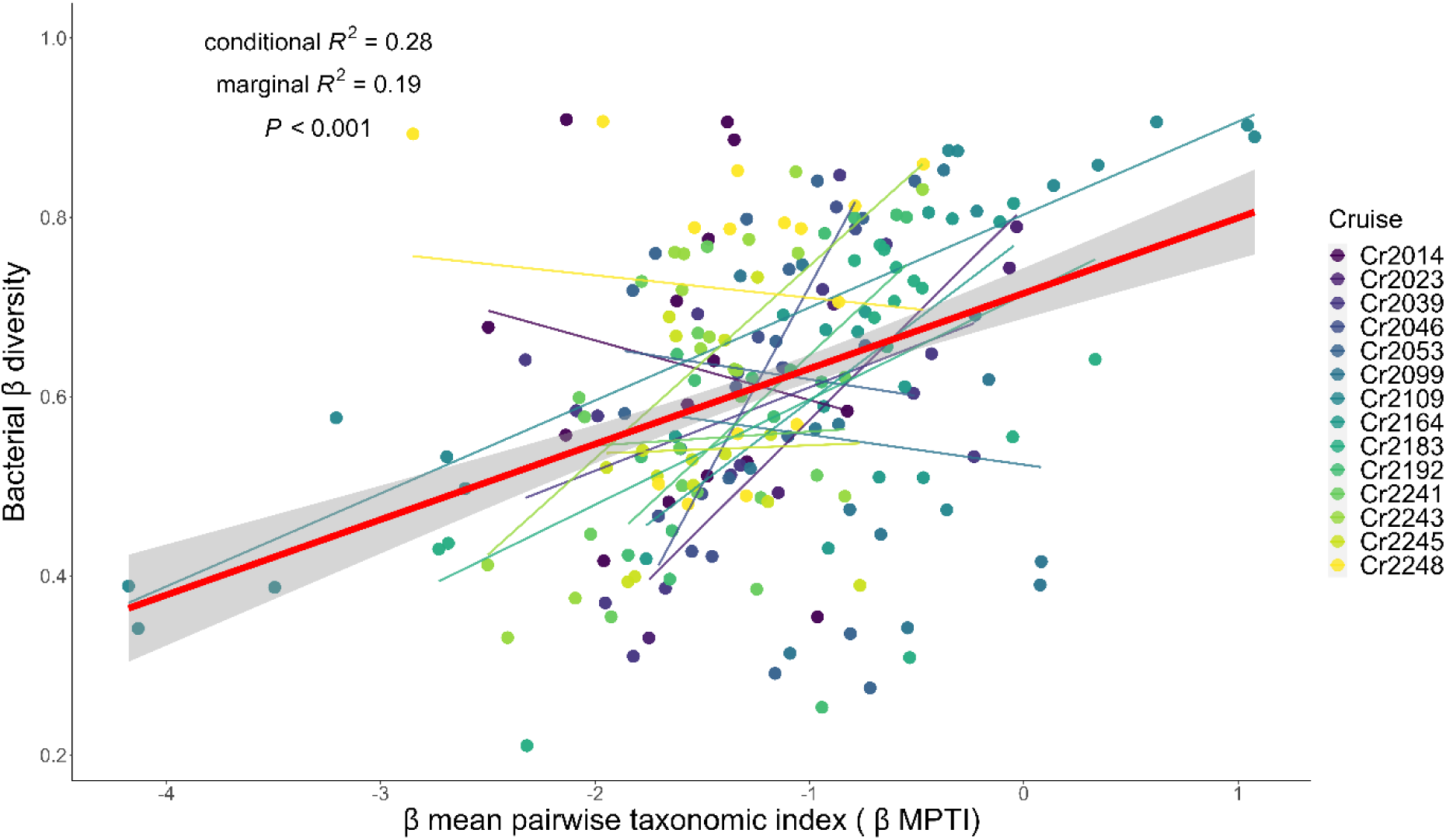
The association between bacterial β diversity and metacommunity assembly processes (*βMPTI*). The bold red solid line indicates the significant regression line fitted by the generalized linear mixed effect model (GLMM), with cruise as the random effect. Grey shaded area represents the confidence interval of regression. The colored dots represent measurements from different sampling cruises. The thin colored solid line represents the regression line fitted by generalized linear model for each sampling cruise.

Through cross-cruise comparison, we further found that when *βMPTI* became less negative, the effects of bacterial β diversity on the biomass of bacterial metacommunity became more positive (Figure 4). This positive relationship remained when statistically controlling for environmental factors (Table 2). Given these results, Hypothesis II is supported, indicating that when non-random/deterministic processes selected for phylogenetically dissimilar species to assemble a metacommunity (less negative *βMPTI*), β diversity was increased and the effects of β diversity on the sum of bacterial communities became more positive.

**Figure 4.**
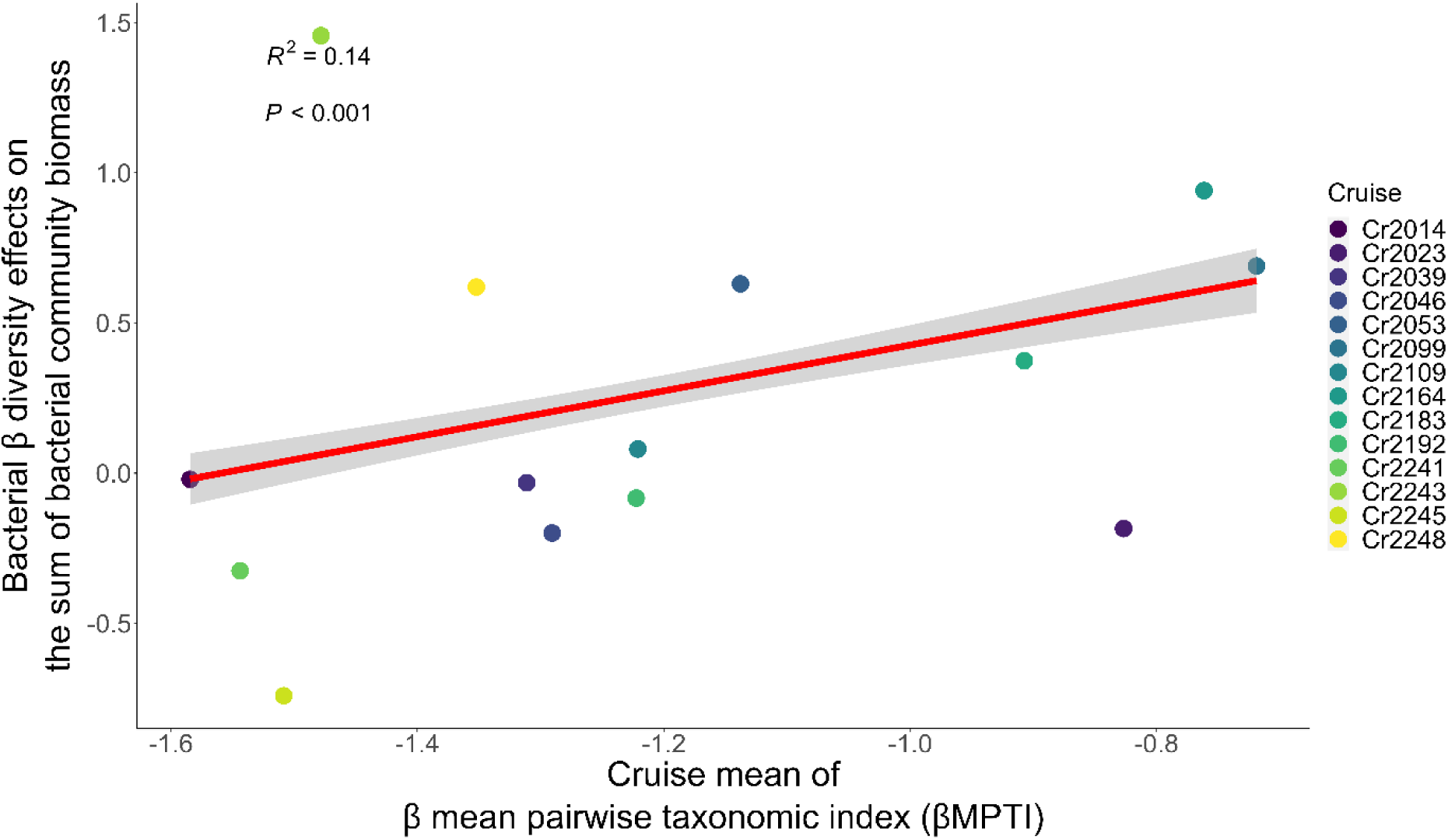
The positive association between bacterial β diversity effects on the sum of bacterial community biomass and metacommunity assembly processes (*βMPTI*) across cruises. The red solid line indicates the significant regression line fitted by the generalized linear model (GLM). Grey shaded area represents the confidence interval of regression. Each of the colored dots represents the slope extracted from the GLM regressing the sum of bacterial community biomass on bacterial β diversity for each of the 14 sampling cruises (c.f. Figure 2).

**Table 2.** Results of generalized linear model (GLM) testing Hypothesis 2: The effect of bacterial β diversity is stronger when β diversity is increased by deterministic assembly processes (Figure 1B). The model presented here is the results after backward stepwise selection to statistically control for potential confounding factors. Steps of the backward stepwise selection is listed in Supplementary Table S4-2. The results indicate that, after accounting for environmental variables, the effect of bacterial β diversity on the summed bacterial community biomass increased with niche-based assembly processes, i.e., non- random/deterministic processes selected for dissimilar species in the metacommunity. This conclusion is qualitatively the same as that drawn from the univariate GLM (Figure 3).

| Bacterial $\beta$ diversity effects on the summed bacterial community biomass as the dependent variable | | | |
| --- | --- | --- | --- |
| Independent variable | Regression coefficient | Standard error | p-value |
| $\beta MPTI$ | 0.79 | 0.06 | <0.01 |
| Log (Salinity) | -22.81 | 1.53 | <0.01 |
| Log (Total inorganic nitrogen) | 0.4 | 0.02 | <0.01 |
| Log (Phosphate) | -0.63 | 0.03 | <0.01 |
| Log (PAR) | -0.05 | 0.02 | <0.01 |
| Log (Chlorophyll-a) | 0.37 | 0.03 | <0.01 |

## Discussion

In the southern ECS, we first confirm that the environment is spatially heterogeneous, and the bacterial composition is thereby affected by spatial heterogeneity (Figure S2-1 and S2-2). Moreover, the degree of spatial heterogeneity varied among cruises. These results are in agreement with previous studies in this region [45, 46, 53]. Given the heterogeneous environments and bacterial composition, we investigate the association between bacterial β diversity and metacommunity functioning. In general, we found a positive, but highly variable, effect of bacterial β diversity on the sum of bacterial communities (Figure 2). The β diversity effects were not significant in 8 out of the 14 cruises (colored dash lines in Figure 2). Current research on the relationship between β diversity and metacommunity functions also find such similar pattern that β diversity generally has positive effects on ecosystem functions [73–77], but non-significant or even negative β diversity effects are not uncommon [8, 9, 78]. Interestingly, we found the variability of β diversity effects coincided with the variability of metacommunity assembly processes, as indicated by *βMPTI* (Figure 3 and S3-1). Accordingly, we argue that the variable β diversity effects may be caused by different strengths of metacommunity assembly processes that determine the β diversity [39].

We first observed that bacterial metacommunity was largely subject to homogenizing assembly processes as most of *βMPTI* was negative (Figure 3). According to the literature, bacterial communities are often found to be governed by homogenizing assembly processes [79, 80], suggesting that bacterial phylogeny is conservative. More importantly, through cross-cruise analysis, we showed that β diversity effects became more positive when β diversity was increased by deterministic assembly processes selecting for phylogenetically dissimilar species (Figure 4). This finding is analogous to the studies revealing that deterministic assembly processes could select for bacteria with certain metabolic capabilities, so that the bacterial communities could utilize resources in the environment; these selected bacterial communities in turn determined the ecosystem functioning [81–83]. Also relevant but to a lesser extent, other studies show that the function of metacommunity is enhanced when deterministic assembly processes select for species that are dissimilar from each other [84, 85]. Together with these studies, our results suggest that when bacterial communities are selected by deterministic, rather than stochastic, assembly processes, they are more capable of capturing resources and performing ecosystem functions. Consequently, when dissimilarity among bacterial communities (β diversity) is increased by deterministic assembly processes, more resources can be utilized so that the effect of β diversity on metacommunity function is further enhanced.

Our findings echo the assertion that community assembly processes can alter the diversity effect on ecosystem functioning [37, 39]. To advocate the influences of assembly processes, previous studies typically show that diversity is maximized by intermediate levels of dispersal that reduce local environmental and demographic stochasticity without imposing too strong homogenization [86–88]. However, whether our study is comparable to this line of research is debatable because dispersal cannot be unarguably defined as a deterministic or stochastic assembly processes even for microbial communities [89, 90]. Since dispersal cannot be easily defined and parsed out [bur see 85, 86], we limit ourselves from making inferences based on dispersal or other specific processes like species interactions. In fact, this is one of the limitations of our analytical framework. Our analyses should be viewed as an examination of collective influences of deterministic versus stochastic assembly processes but not an examination of certain specific ecological processes. Future studies dedicated to examining the influences of specific assembly processes are encouraged and should render fruitful results.

Some caveats associated with our analytical framework is worth mentioning. First, we are aware that inter-dependency among α, β, and γ diversity could lead to statistical artifacts [20, 21]. We did find that bacterial β diversity was correlated with the richness and Shannon diversity of the metacommunity (correlation coefficient = 0.14 and 0.18 respectively; both p-values < 0.01; Figure S4-1). The richness and Shannon diversity was calculated from the combined communities of any pair of the 6 stations in the same sampling cruise (metacommunity). However, the summed bacterial community biomass was not significantly associated with the richness and Shannon diversity (Figure S4-2). We thus argue that our conclusions are robust to those statistical artifacts.

Moreover, we are aware that the bacterial biomass we measured was a merely proxy because it is calculated by the multiplication of cell count and conversion factor. Nevertheless, such conversion is widely acceptable [93]. We also acknowledge that biomass is not a perfect measurement of ecosystem functioning although it has been commonly used [17, 94–96]. Future studies applying metagenomic or metatranscriptomic analyses to our analytical framework will render a direct test of biodiversity-ecosystem functioning relationship.

Next, our findings hinge on the assumption that *βMPTI*, or phylogenetic similarity in general, can be used to imply the occurrence of community assembly processes. Because *βMPTI* is calculated in the similar fashion as the net related index (NTI), nearest taxon index (NRI) [41, 66], βNTI, and βNRI [67], we argue that *βMPTI* could indicate whether the community is subject to certain assembly processes such as environmental filtering [97] or competition [98]. The βNTI and βNRI have recently been used to indicate the assembly processes that determine the composition of microbial community [29, 79, 80]. We also observed significant phylogenetic signals (Figure S5-1), which demonstrated the relationship between phylogenetic relatedness and ecological similarity [42]. However, we acknowledge that phylogenetic similarity might not faithfully approximate community assembly processes [99, 100]. We thus cautiously treat the *βMPTI* as an implication of the occurrence of certain community assembly processes.

Despite these limitations, our analytical framework expands the current biodiversity-ecosystem functioning framework from a local (α) level to a metacommunity (β) level. Because natural ecological communities are inherently interconnected, expanding to the metacommunity level allows us to holistically perceive the role of biodiversity in natural ecosystems. In addition, we demonstrate how the community assembly processes alter the effects of β diversity on the functioning of metacommunity. Taking the influences of community assembly processes into account equips us a more mechanistic understanding of the impacts of biodiversity on ecosystem functioning.

In conclusion, we showed that bacterial β diversity positively affected the summed bacterial community biomass in the southern ECS. Moreover, this positive effect was stronger when bacterial β diversity was increased by the assembly processes that deterministically (i.e., non-randomly) selected for phylogenetically dissimilar species. Our findings support that, in the metacommunity (β) level, niche-based deterministic community processes further enhance the effect of biodiversity on ecosystem functioning of bacterial community in the southern ECS.

## Acknowledgements

We thank Hon-Tsen Yu for providing facilities and advice on laboratory work. We also thank Sara Jackrel for her feedbacks on the manuscript. This work was supported by the National Center for Theoretical Sciences, Foundation for the Advancement of Outstanding Scholarship, and the Ministry of Science and Technology, Taiwan.

## Data Availability Statement

The sequence data have been deposited in the NCBI Sequence Read Archive (SRA) under the accession numbers: PRJNA662424 (https://www.ncbi.nlm.nih.gov/bioproject/?term=PRJNA662424).

## Supplement 1

### 1.1 Library preparation and sequencing

Total DNA was extracted separately from the 0.2 μm-pore size filters with the PowerWater DNA Extraction Kit (Qiagen) according to the manufacturer’s instructions. DNA extracts from the 0.2-μm-pore size filters were used as templates of polymerase chain reaction (PCR) for 16S sequencing. PCR was performed in two steps: to amplify targeted 16S regions in the first step PCR, and attach sample-specific barcodes for each sample in the second step PCR [1].

For the prokaryotic 16S rDNA, the V5–V6 region of 16S rDNA was amplified using the forward primer FIA-787F (5’-[illumina forward index adaptor]-ATTAGATACCCNGGTAG- 3’) and reverse primer RIA-1046R (5’-[illumina reverse index adaptor]- CGACAGCCATGCANCACCT-3’) [2]. In the first step PCR, 20-μl of PCR mix contained: 1 U of Taq DNA polymerase (Promega), 1 × reaction buffer, 1.5 mM MgCl2, 0.2 mM dNTPs, 0.2 mM of primers and 2 ng DNA templates. Conditions for the PCR cycles were: an initial denaturation at 94 °C for 3 min; 25 cycles of 94 °C for 30 s, 55 °C for 45 s, 72 °C for 1 min; and a final extension at 72 °C for 2 min. Three PCR replicates were conducted and pooled for each sample for obtaining enough DNA concentration for sequencing.

DNA products of the first step PCR were purified using AMPure XP beads (Beckman Coulter Genomic, CA, USA) and DNA concentration was quantified using a Qubit fluorometer (Invitrogen, Carlsbad, CA, USA) with Qubit dsDNA BR Assay Kit (Life Technologies, USA) before conducting the second step PCR.

Primers containing sample-specific barcodes and Illumina adaptors were used to perform the second step PCR. 20-μl of PCR mix contained: 1 U of Taq DNA polymerase (Promega), 1 × reaction buffer, 1.5 mM MgCl2, 0.2 mM dNTPs, 0.2 mM of S5, and N7 primers (Nextera Index Kit) and 2 ng DNA obtained from the first step PCR. The PCR conditions was with initial denaturation at 94 °C for 3 min; 6 cycles of 94 °C for 30 s, 55 °C for 45 s, 72 °C for 1 min; and a final extension at 72 °C for 2 min. For this, three PCR replicates were conducted and pooled for each sample. Finally, DNA samples with unique barcodes were pooled in roughly equal concentrations and sent for 2 × 300 bp paired-end sequencing with Illumina Miseq platform. The sequence data have been deposited in the NCBI Sequence Read Archive (SRA) under the accession numbers: PRJNA662424.

### 1.2 Sequence processing

Sequences were analyzed using DADA2 v.1.12 [3] pipeline running on R v.3.6.2. Suggested by DADA2 pipeline tutorial, paired-end sequences were filtered and trimmed separately depends on sequences quality by using function “filterANDTrim” before DADA2 algorism. Primer sequences were trimmed, and sequences were truncated at position 220-bp (for forward reads) and 180-bp (for reverse reads) for 16S rDNA sequences, where average quality score start to crash lower than Q30. We thus include the sequences that have average quality score greater than Q30. Sequences contain Ns and more than two expected errors were also removed. DADA2 algorithm was then performed to remove spurious sequences that possibly be generated during PCR amplification and sequencing [3]. After quality filtration, paired-end reads were merged using function “mergePairs” with a minimum of 12- bp overlap. Merged sequences were then assembled in to ASVs (amplicon sequence variant) with function “makeSequenceTable”. Finally, chimeras were detected and removed from ASV pool using “removeBimeraDenovo”.

Taxonomy assignment was preformed to recognize bacterial ASVs from 16S rDNA. Six taxonomy levels (from phylum to the genus) were assigned using Naïve Bayesian Classifier [4] implemented in function “assignTaxonomy”. The exact matching to the species level was assigned using function “assignSpecies”. For reference databases, Silva 132 database [5] was used for assigning 16S rDNA reads. Subsequently, bacterial communities were selected as ASVs classified under kingdom “bacteria”.

With the aim to quantify bacteria community assembly processes with phylogeny information, phylogenetic trees were built with maximum likelihood method using the “phangorn” package in R (with negative edges length = 0).

## Supplement 2

**Figure S2-1.**
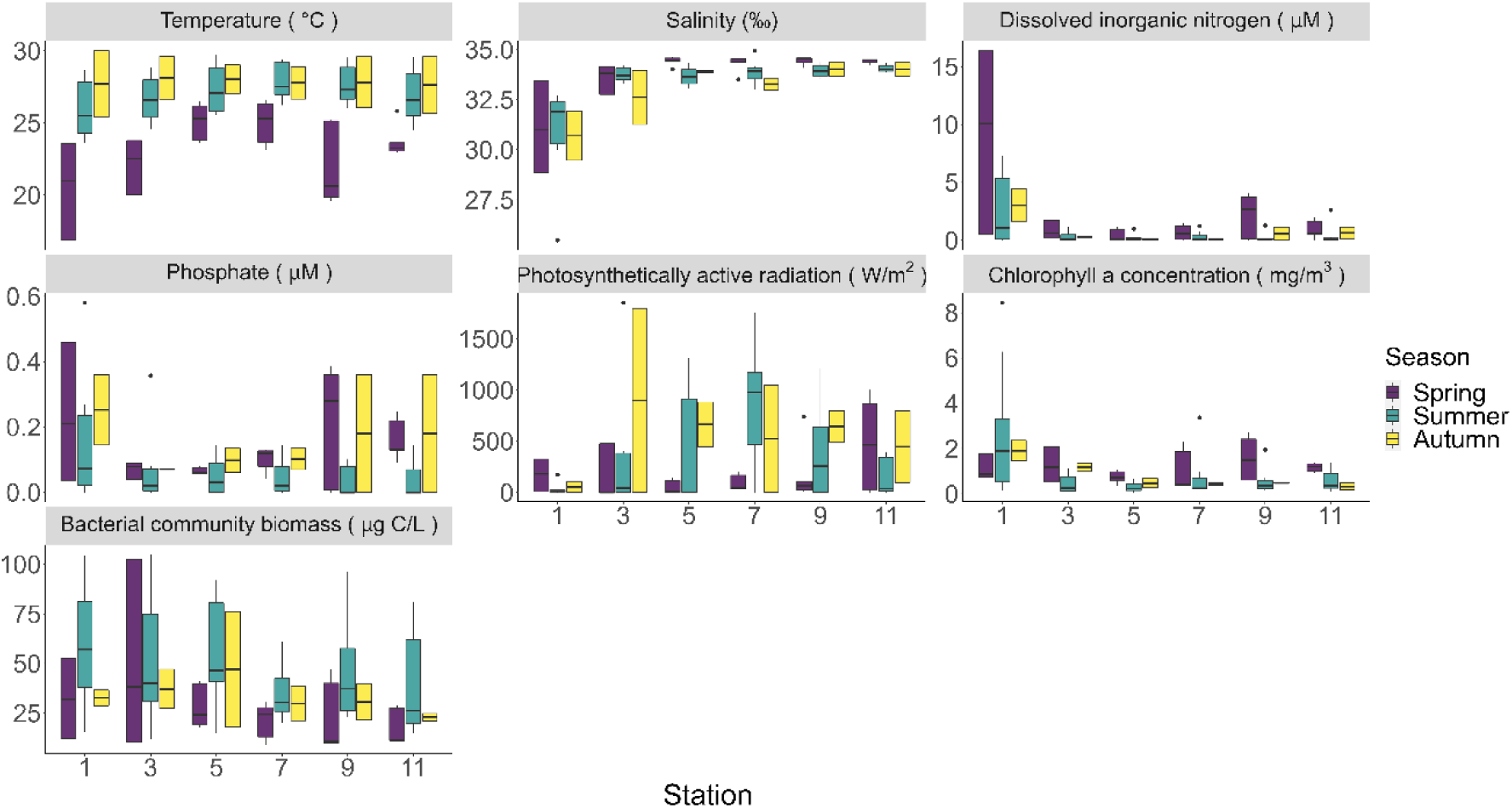
Box plots of environmental variables and bacterial community biomass of the 6 sampling stations in the southern East China Sea. The color represents the season of the cruise (spring is defined as March, April, and May; summer is defined as June, July, and August; Autumn is defined as September, October, and November).

**Figure S2-2.**
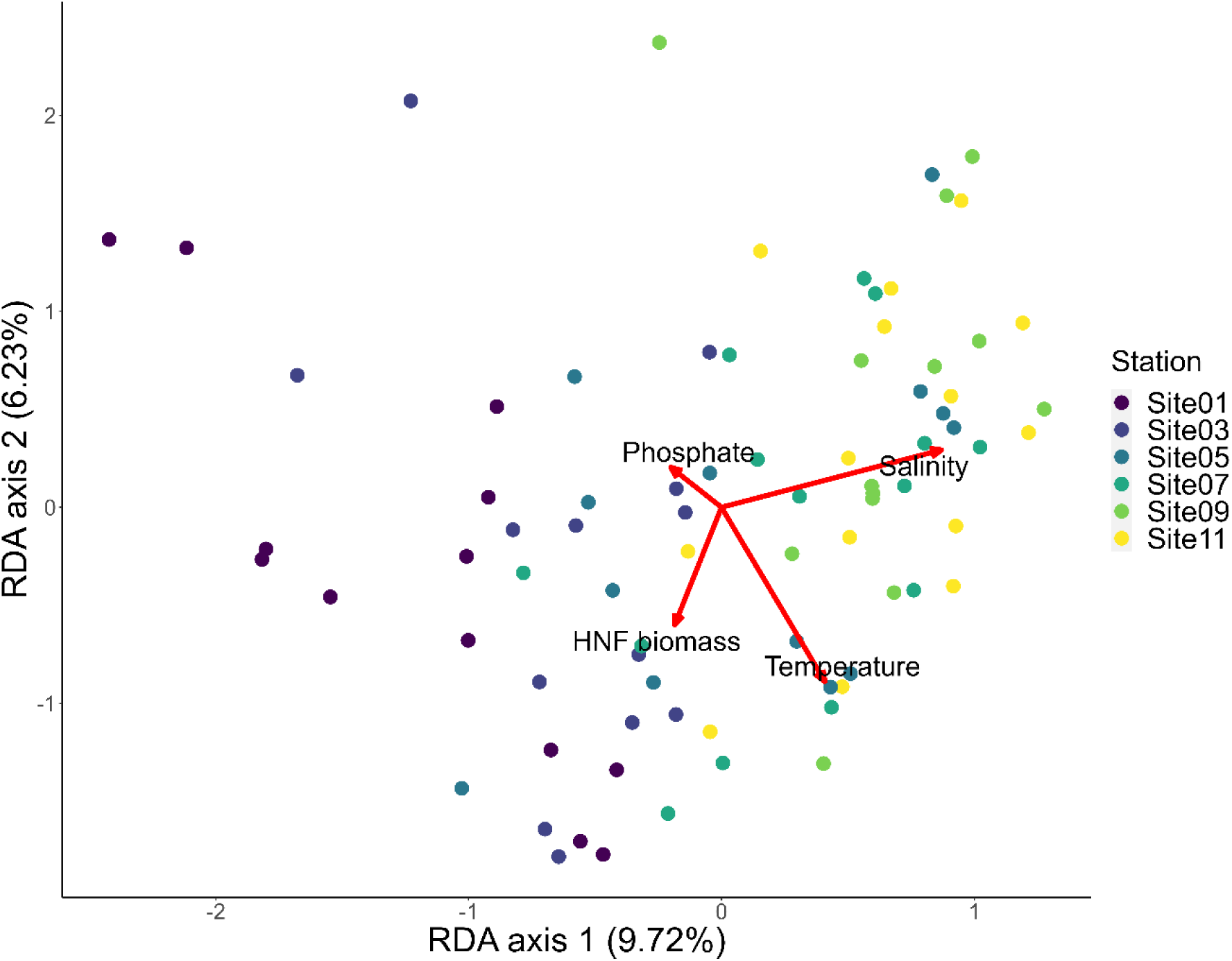
Biplot of distance-based redundancy analysis showing the relationships between bacterial taxonomic composition and environmental variables. Dissimilarities among bacterial communities are the Bray-Curtis dissimilarities between bacterial communities in the same cruise. The red arrows represent the environmental variables that significantly explain the dissimilarities among bacterial communities. HNF biomass (μg C/L) is the community biomass of heterotrophic nanoflagellates (HNF), which are the potential predators of bacteria. The colors of dots represent different sampling stations.

## Supplement 3

**Supplementary Table S3-1.** The backward selection sequences to test Hypothesis I: Bacterial Bray-Cutis dissimilarity index increases the sum of bacterial community biomass (Figure 1B). The bacterial β diversity is always included in each step. The variable labeled with an asterisk is the factor with the highest p-value in each step and is removed in the next step.

| Summed bacterial community biomass as the dependent variable |  |  |
| --- | --- | --- |
|  | Independent variable | p-value |
| Selection step 1 |  |  |
| | Log (Bacterial $\beta$ diversity) | 0.02 |
|  | Log (Temperature) | 0.17 |
|  | Log (Salinity) | <0.01 |
|  | Log (Total inorganic nitrogen) | <0.01 |
|  | *Log (Phosphate) | 0.77 |
|  | Log (PAR) | 0.11 |
|  | Log (Chlorophyll-a) | 0.31 |
| Selection step 2 |  |  |
| | Log (Bacterial $\beta$ diversity) | 0.02 |
|  | Log (Temperature) | 0.17 |
|  | Log (Salinity) | <0.01 |
|  | Log (Total inorganic nitrogen) | <0.01 |
|  | Log (PAR) | 0.11 |
|  | *Log (Chlorophyll-a) | 0.33 |
| Selection step 3 |  |  |
| | Log (Bacterial $\beta$ diversity) | 0.02 |
|  | *Log (Temperature) | 0.16 |
|  | Log (Salinity) | <0.01 |
|  | Log (Total inorganic nitrogen) | <0.01 |
|  | Log (PAR) | 0.14 |
| Selection step 4 |  |  |
| | Log (Bacterial $\beta$ diversity) | 0.03 |
|  | Log (Salinity) | <0.01 |
|  | Log (Total inorganic nitrogen) | <0.01 |
|  | Log (PAR) | 0.17 |
| Selection step 4 |  |  |
| | Log (Bacterial $\beta$ diversity) | 0.04 |
|  | Log (Salinity) | <0.01 |
|  | Log (Total inorganic nitrogen) | <0.01 |

**Supplementary Table S3-2.** The backward selection sequences to test Hypothesis II: Effects of β diversity on the summed bacterial community biomass are increased when niche-based assembly processes (i.e., non-random/deterministic processes) select for dissimilar species (Figure 1C). The *βMPTI* is always included in each step. The variable labeled with an asterisk is the factor with the highest p-value in each step and is removed in the next step.

|  | Independent variable | p-value |
| --- | --- | --- |
| Selection step 1 |  |  |
| | $\beta MPTI$ | <0.01 |
|  | *Log (Temperature) | 0.22 |
|  | Log (Salinity) | <0.01 |
|  | Log (Total inorganic nitrogen) | <0.01 |
|  | Log (Phosphate) | <0.01 |
|  | Log (PAR) | 0.01 |
|  | Log (Chlorophyll-a concentration) | <0.01 |
| Selection step 2 |  |  |
| | $\beta MPTI$ | <0.01 |
|  | Log (Salinity) | <0.01 |
|  | Log (Total inorganic nitrogen) | <0.01 |
|  | Log (Phosphate) | <0.01 |
|  | Log (PAR) | 0.01 |
|  | Log (Chlorophyll-a) | <0.01 |

**Figure S3-1.**
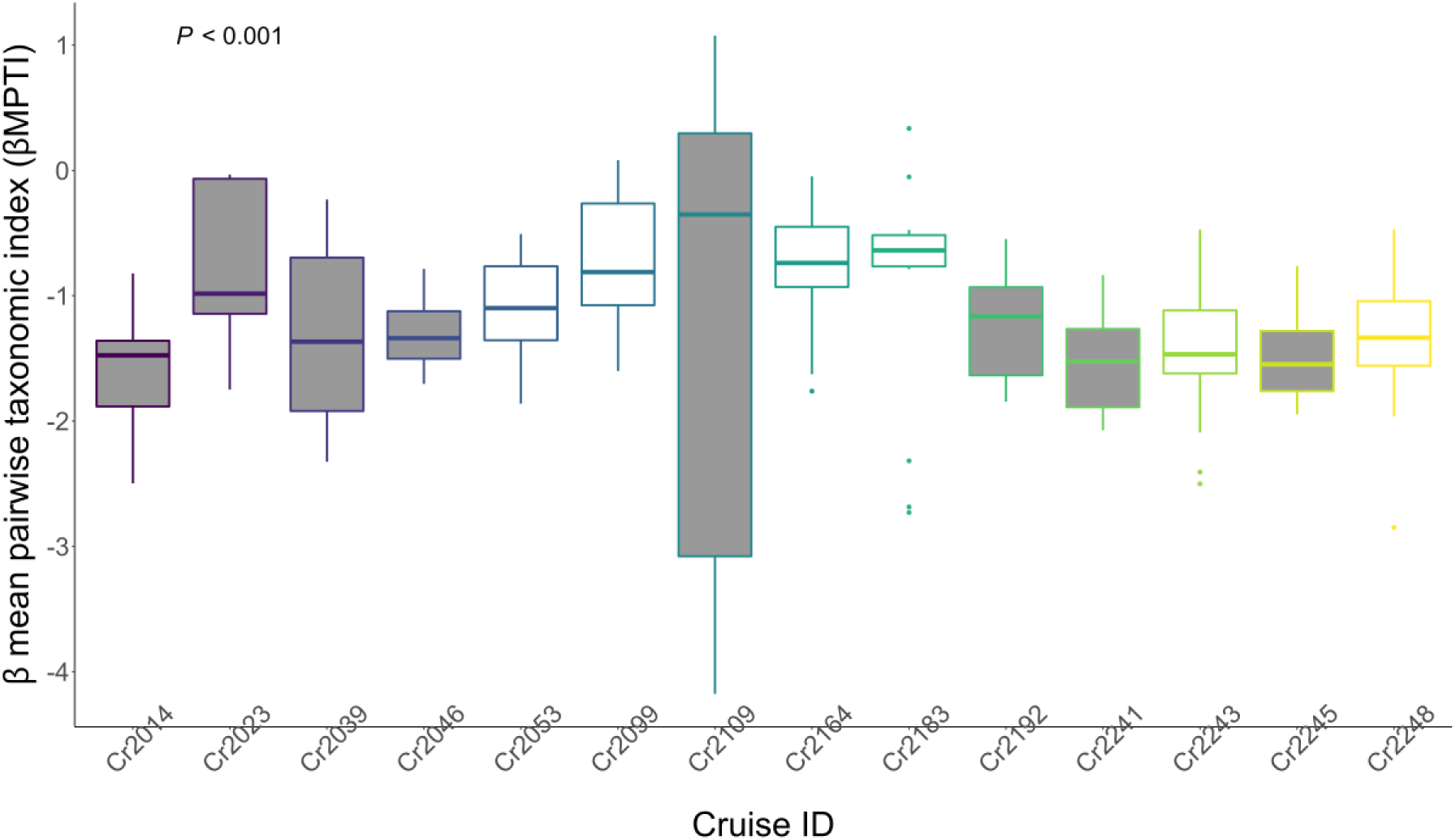
Box plot showing the variation of *βMPTI* among the 14 sampling cruises. The colors represent different sampling cruises. The grey filled boxes are the cruises in which the association between β diversity and the summed bacterial community biomass is negative or non-significant.

## Supplement 4

**Figure S4-1.**
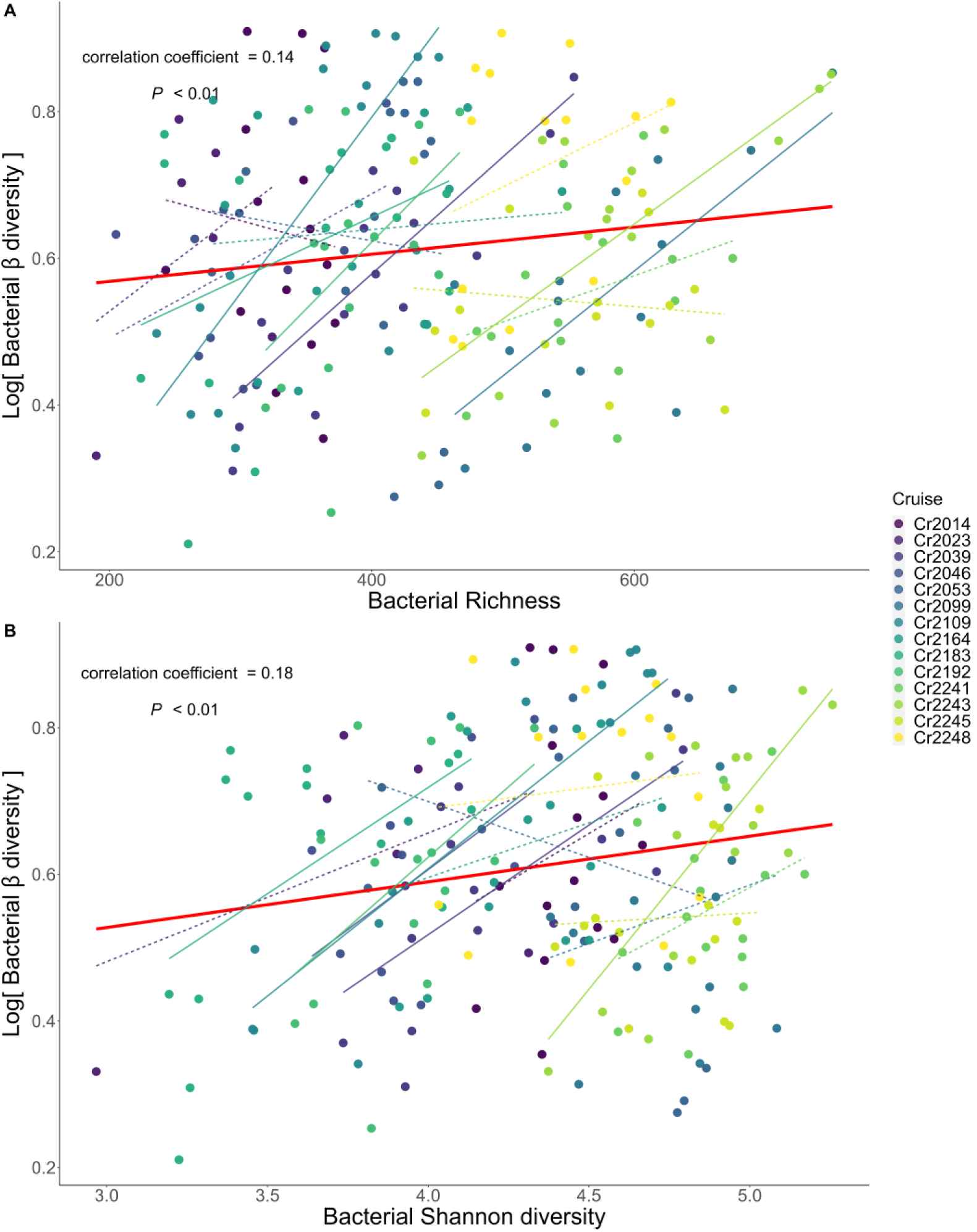
The correlation between β diversity versus pooled richness (A) and Shannon diversity (B) of the bacterial metacommunity. The richness and Shannon diversity was calculated from the combined community composition of the two communities comprising the metacommunity. The red solid line indicates the significant positive correlation. The colored dots represent measurements from different sampling cruises. The thin colored line represents the regression lines for each cruise. The colored solid and dashed lines represent significant and non-significant correlations, respectively.

**Figure S4-2.**
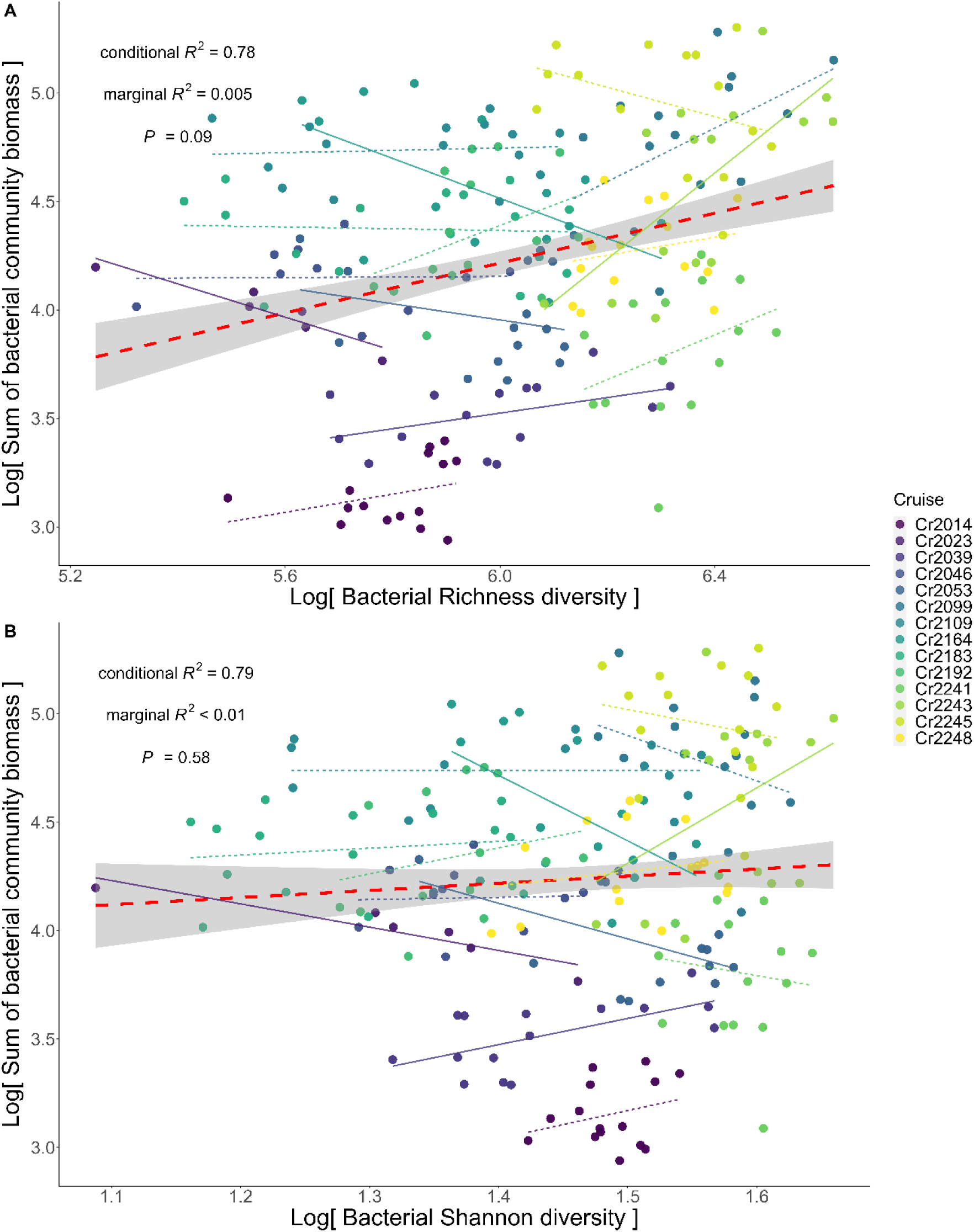
The non-significant association between the summed bacterial community biomass versus pooled richness (A) and Shannon diversity (B) of the bacterial pair-wise metacommunity. The richness and Shannon diversity was calculated from the combined community composition of any pairs of the 6 stations in the same sampling cruise. The red dashed line indicates the regression line fitted by the generalized linear mixed effect model (GLMM), with cruise as the random effect. Grey shaded area represents the confidence interval of regression. The colored dots represent measurements from different sampling cruises. The thin colored line represents the regression line fitted by generalized linear model regressing metacommunity biomass on pooled richness (A) and Shannon diversity (B) for each cruise. Colored solid and dashed lines represent significant and non-significant regression respectively.

## Supplement 5

In this supplement, we show the phylogenetic signal for the bacteria community in the southern East China Sea. To do so, we first quantified the abiotic habitats for each ASV (amplicon sequence variant) by calculating the abundance-weighted average of environmental variables (including temperature, salinity, photosynthetic active radiation, nitrite, nitrate and phosphate concentrations). We then plotted the Mantel correlations between ASVs’ habitats against the phylogenetic distance among all ASVs (Figure S5-1). The phylogenetic signal was detected using the *mantel.correlog* function in the vegan package in R4.0.0. Significant phylogenetic signal suggests that phylogenetic distances can be used to approximate the ecological niche of species [6]

**Figure S5-1.**
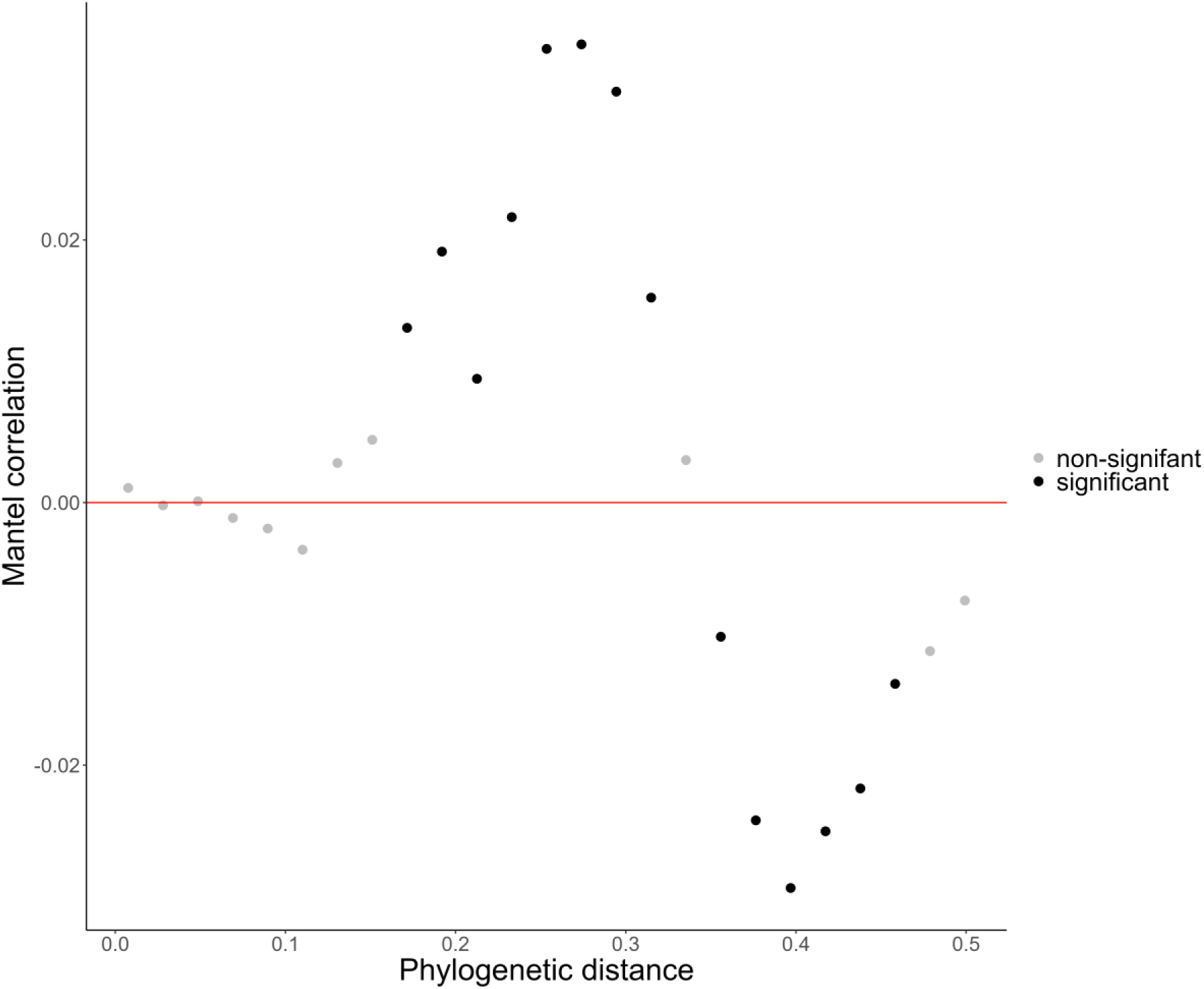
Phylogenetic Mantel correlogram showing significant phylogenetic signals of habitat preferences in bacteria community. Black and grey dots denote significant and nonsignificant correlations, respectively, relating between-ASV habitat differences to between-ASV phylogenetic distances.

